# Resistance-guided mining of bacterial genotoxins defines a family of DNA glycosylases

**DOI:** 10.1101/2021.10.29.466481

**Authors:** Noah P. Bradley, Katherine L. Wahl, Jacob L. Steenwyk, Antonis Rokas, Brandt F. Eichman

## Abstract

Unique DNA repair enzymes that provide self-resistance against genotoxic natural products have been discovered recently in bacterial biosynthetic gene clusters (BGCs). The DNA glycosylase AlkZ belongs to a superfamily of uncharacterized proteins found in antibiotic producers and pathogens, but despite its importance to azinomycin B resistance, the roles of AlkZ orthologs in production of other natural products are unknown. Here, we analyze the genomic distribution and use a resistance-based genome mining approach to identify *Streptomyces* AlkZ homologs associated with known and uncharacterized BGCs. We show that the ortholog associated with synthesis of the alkylating agent hedamycin excises hedamycin-DNA adducts and provides resistance to the genotoxin in cells. Our results define AlkZ in self-resistance to specific antimicrobials and implicate a related but distinct homolog, which we name AlkX, in protection against an array of genotoxins. This work provides a framework for targeted discovery of new genotoxic compounds with therapeutic potential.

## Introduction

Bacteria are exceptionally rich sources of secondary metabolites, which are important for their survival and often have therapeutic value. *Streptomyces* produce 35% of all known microbial natural products and nearly 70% of all commercially useful antibiotics, with several being FDA-approved antitumor agents used as first-line cancer treatments (Demain and Sanchez, 2009; Jacob and Weissman, 2017; Law et al., 2020; Procopio et al., 2012). Secondary metabolites are often toxins used in ecological interactions with other organisms and can target any number of critical cellular functions (Seyedsayamdost, 2019; Tyc et al., 2017). Natural products that damage DNA (genotoxins) form covalent or non-covalent DNA adducts that can inhibit replication and transcription, thus undermining genomic integrity through mutagenesis or cell death (Chumduri et al., 2016; Gates, 2009). Consequently, genotoxins are particularly useful antineoplastic agents, as exemplified by several clinically relevant drugs including doxorubicin, bleomycin, mitomycin C, and duocarmycin analogs (Huang et al., 2021).

*Streptomyces* produce a wide variety of DNA alkylating and oxidizing agents that have antimicrobial and antitumor properties. Spirocyclopropylcyclohexadienones (duocarmycin A and SA, yatakemycin, CC-1065) (Boger and Garbaccio, 1997; Parrish et al., 2003), pluramycins (pluramycin A, hedamycin, altromycin) (Hansen and Hurley, 1995; Hansen et al., 1995; Kwok et al., 1998), anthracycline glycosides (trioxacarcin A, LL-D49194α1) (Dong et al., 2019; Pfoh et al., 2008), and the leinamycin family (Nooner et al., 2004) contain a single reactive group that covalently modifies purine nucleobases to form a broad spectrum of bulky alkyl-DNA monoadducts. *Streptomyces* also produce bifunctional alkylating agents that react with nucleobases on both DNA strands to create interstrand crosslinks (ICLs). Mitomycin C (MMC) from *S. lavendulae* crosslinks guanines at their N2 positions, and azinomycin A and B (AZA, AZB) from *S. sahachiroi* and *griseofuscus* crosslink purines at their N7 nitrogens (Bizanek et al., 1992; Coleman et al., 2002). In addition to alkylating agents, several families of natural products, including bleomycins and enediynes, exert their toxicity by oxidative cleavage of DNA and RNA (Galm et al., 2005).

The production of secondary metabolites in *Streptomyces* is genetically organized into biosynthetic gene clusters (BGCs), which contain the genes necessary for their biosynthesis, export, regulation, and resistance. Resistance mechanisms protect antibiotic producers from toxicity of their own natural products, and include toxin sequestration, efflux, modification, destruction, and target repair/protection (Cundliffe and Demain, 2010; D’Costa et al., 2006; Tenconi and Rigali, 2018). In the case of genotoxins, several DNA repair enzymes have been identified as target repair resistance mechanisms, including direct reversal of streptozotocin alkylation by AlkB and AGT (alkylguanine alkyltransferase) homologs (Ng et al., 2019), base excision of yatakemycin-adenine adducts by the DNA glycosylase YtkR2 (Xu et al., 2012), nucleotide excision of DNA adducts of several intercalating agents including daunorubicin (Furuya and Hutchinson, 1998; Lomovskaya et al., 1996; Prija et al., 2017), and putative replication-coupled repair of distamycin-DNA adducts (Ma et al., 2020).

The AZB BGC in *Streptomyces sahachiroi* encodes a DNA glycosylase, AlkZ, that unhooks AZB-ICLs and that provides cellular resistance against AZB toxicity (Mullins et al., 2019; Wang et al., 2016). ICL unhooking by AlkZ involves hydrolysis of the N-glycosidic bonds of the crosslinked deoxyguanosine residues, producing abasic (AP) sites that can be repaired by the base excision repair pathway (Mullins et al., 2019). AlkZ belongs to the relatively uncharacterized HTH_42 superfamily of proteins found largely in antibiotic-producing and pathogenic bacteria (Wang et al., 2016). The crystal structure of AlkZ revealed a unique C-shaped architecture formed by three tandem winged helix-turn-helix motifs, with two glutamine residues (QΦQ motif) essential for catalysis located at the center of the concave surface (Mullins et al., 2017). We recently characterized a second HTH_42 protein from *Escherichia coli*, YcaQ, as a DNA glycosylase within a four-gene operon that excises several types of guanine N7-linked ICLs and provides *E. coli* with resistance to the nitrogen mustard, mechlorethamine (Bradley et al., 2020). In addition to ICLs, both AlkZ and YcaQ are able to excise N7-methylguanine (7mG) monoadducts from DNA (Bradley et al., 2020). Henceforth, we will refer to YcaQ as AlkX (X-link/crosslink) to reflect its role as an alkylation specific ICL repair enzyme.

The targeted discovery of natural products has been employed to search for novel scaffolds in plants, fungi, and bacteria, and can be useful for identifying specific classes of compounds (Belknap et al., 2020; Kersten and Weng, 2018; Kjaerbolling et al., 2019; Shigdel et al., 2020). Genome mining can be used to search for unidentified BGCs through analysis of core/accessory biosynthetic genes (PKS, NRPS, tailoring enzymes), comparative/phylogeny-based mining, regulatory genes, and more recently, resistance genes (Ziemert et al., 2016). Some of these resistance-based mining approaches focus on the experimental screening of antibiotic resistance, while others rely on bioinformatic tools to identify resistance genes within clusters based on homology to known resistance genes (Blin et al., 2019; Mungan et al., 2020; Skinnider et al., 2015; Thaker et al., 2013). However, many of these resistance-based methods have not been applied in bacteria for targeted discovery.

Here, we characterize the genomic differences between AlkX and AlkZ glycosylases found in 435 species of *Streptomyces* to develop additional insight into this new family of DNA repair proteins, and apply this information in resistance-guided genome mining to characterize unknown BGCs or identify new genotoxins. We found that AlkZ homologs are highly variable in sequence and enriched in BGCs, many producing known genotoxic alkylating agents. We show that the AlkZ homolog within the BGC of a known DNA alkylating agent, hedamycin, is a resistance DNA glycosylase specific for hedamycin-guanine lesions, consistent with AlkZ-mediated DNA repair activity as a general self-resistance mechanism to genotoxins in antibiotic producers. Moreover, we found AlkZ homologs in BCGs that are either uncharacterized or that produce natural products not previously known to be genotoxic, validating resistance genome mining as an approach to discover new genotoxins. We also found that AlkX homologs are highly conserved in sequence and genetic neighborhood and are not associated with BGCs, suggesting a broader role of HTH_42 superfamily proteins outside of antibiotic self-resistance in bacteria.

## Results

### AlkX and AlkZ homologs in Streptomyces are evolutionarily distinct

*E. coli* AlkX and *S. sahachiroi* AlkZ are the only characterized members of the HTH_42 superfamily and are unique in their ability to unhook ICLs and to provide cellular resistance to crosslinking agents. Both enzymes fully unhook ICLs derived from AZB (Fig. 1A). While AlkZ is specific for AZB-ICLs and is essential to the AZB-producing organism, AlkX has a relaxed specificity and can unhook ICLs derived from the simple bifunctional alkylating agent mechlorethamine (Fig. 1B) (Bradley et al., 2020; Wang et al., 2016). AlkX and AlkZ belong to one of five classes of HTH_42 proteins characterized by domain organization, which accounts for >95% of all HTH_42 proteins (Fig. S1A). Approximately two-thirds of the known HTH_42 proteins in prokaryotes are found in Actinobacteria, with ~ 25% of those sequences from *Streptomycetales* (Fig. S1B,C). The remainder are found in several different orders of Bacteria, and a very small number (12) in Archaea.

**Figure 1.**
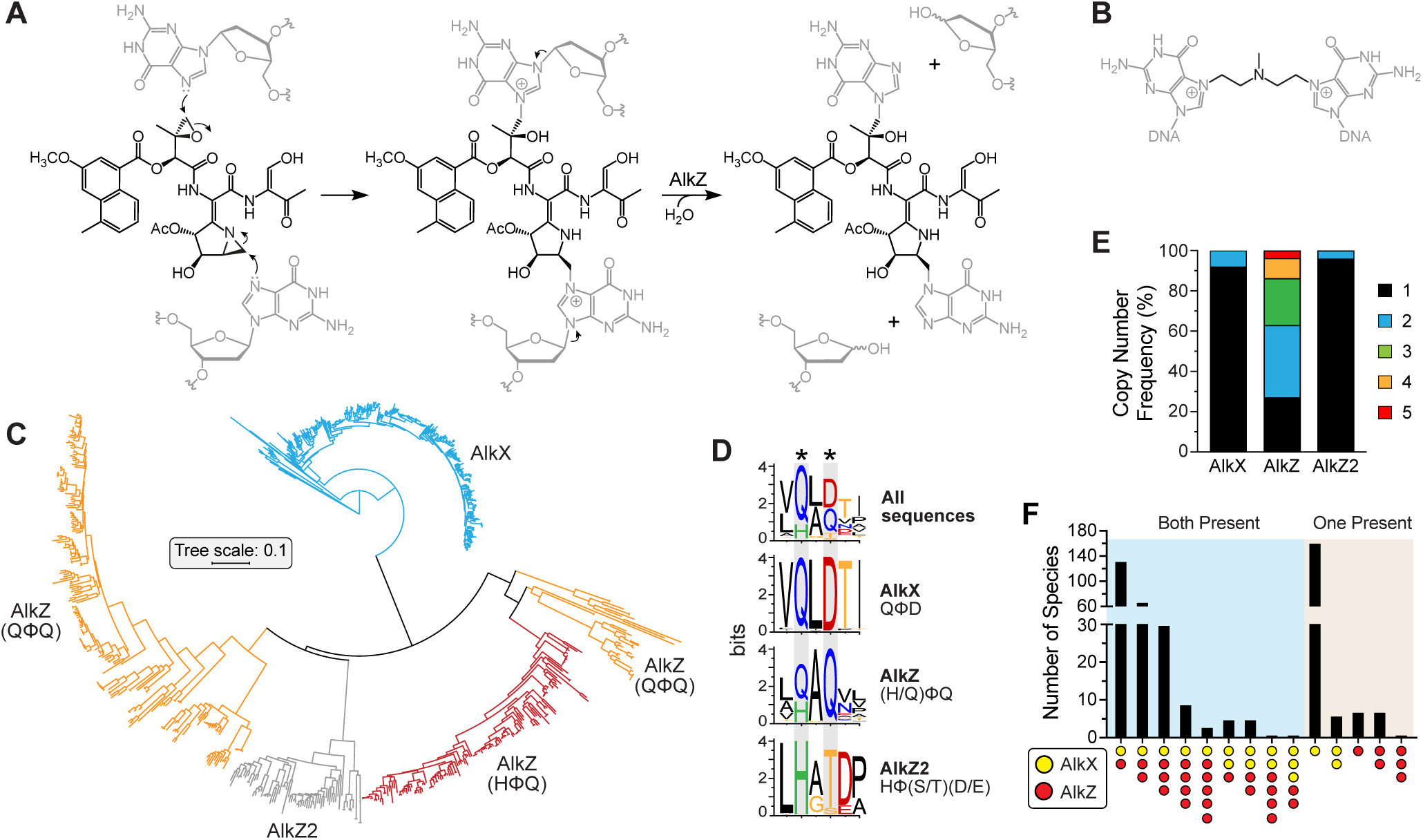
Phylogenetic Organization of AlkX/AlkZ Homologs in *Streptomyces*. (A) Azinomycin B reacts with opposite strands of DNA to form an ICL, which is unhooked by AlkZ-catalyzed hydrolysis. (B) Structure of a nitrogen mustard ICL derived from mechlorethamine and unhooked by AlkX. (C) Phylogenetic tree of *Streptomyces* AlkX/AlkZ homologs (n=897) colored by homolog (AlkX, blue; AlkZ, red/orange; AlkZ2, grey). (D) Sequence logos for the catalytic motifs within *Streptomyces* AlkX and AlkZ homologs. Catalytic residues are marked with asterisks. Colors correspond to side chain chemistry. (E) AlkX/AlkZ copy number frequency per *Streptomyces* genome, as a percentage of the total species analyzed (n=436 species, 897 sequences). One-way ANOVA significance (P) values of copy number variance are 0.0078 (AlkX-AlkZ), 0.0033 (AlkZ-AlkZ2), and 0.3305 (AlkX-AlkZ2), the latter of which is not significant. (F) AlkX/AlkZ coincidence frequency. Blue shaded section represents species containing both AlkX and AlkZ; tan shaded section represents species containing either AlkX or AlkZ.

To better understand the evolutionary and phylogenetic breadth of this superfamily in *Streptomyces*, we collected and analyzed all AlkX and AlkZ protein sequences from available genomes using a combination of BLAST searches against *Streptomyces* genomes in GenBank and HHMR protein domain searches of the BLAST hits against the Pfam database (Table S1). AlkX and AlkZ represent 49% and 43% of the total number of sequences, respectively. Alignment of the 897 sequences showed that AlkX and AlkZ homologs fall into distinct clades that represent (Fig. 1C). The clades are defined in part by unique catalytic motifs QΦD (AlkX) and (Q/H)Φ Q (AlkZ), where Φ is an aliphatic residue (Bradley et al., 2020; Mullins et al., 2017). AlkX homologs show a high degree (>75%) of amino acid sequence conservation, whereas the AlkZ subfamily is more diverse, with only around 40% amino acid similarity on average. The differences in conservation are consistent with mutation rates as approximated by tip-to-root branch lengths (0.23 for AlkX and 0.59 for AlkZ). In addition, we found that 8% of sequences do not fall into either AlkX or AlkZ clades and contain a unique catalytic consensus sequence, HΦ(S/T)(D/E) (Fig. 1C,D). Because these sequences exhibit greater similarity overall to AlkZ than AlkX, we have named this third homolog AlkZ2. However, AlkZ2 is somewhat of a hybrid between AlkZ and AlkX in that it is more similar to AlkX in its copy number and genomic location (see below). We verified that proteins within the AlkZ2 clade contains bona fide DNA glycosylase activity, as *S. caeruleatus* AlkZ2 excised 7mG from DNA in a similar manner to *S. sahachiroi* AlkZ (Fig. S1E).

We also found that AlkZ genes are often found in multiple copies and in different combinations in many species of *Streptomyces*. The copy number differences between the different clades are significant, with the majority (90-95%) of AlkX and AlkZ2 homologs found as a single copy and AlkZ mainly found in multiple (2-5) copies (Fig. 1E). The coincidence of AlkX and AlkZ in the species examined also varies. Although the most common combination is the presence of a copy of each AlkX and AlkZ, many other combinations are observed (Fig. 1F). The number of species that contain both genes decreases as the copy number increases. For species containing either AlkX or AlkZ (not both), the majority contain a single AlkX copy, with just a few species having only AlkZ present. These results show that both AlkX and AlkZ homologs are broadly distributed across *Streptomyces* and are distinct with respect to sequence, diversity, and copy number.

### AlkZ homologs are prevalent in biosynthetic gene clusters

Given the distinct phylogeny of AlkX/AlkZ homologs, we next examined their proximity to BGCs and characterized the identities of clusters containing a homolog. To perform this analysis, we identified all BGCs in the genomes of known *Streptomyces* species containing AlkX/AlkZ homologs, determined the most similar known cluster via BLAST, and extracted the distance in base pairs between the AlkX/AlkZ homolog and the nearest 3’ or 5’ end of each BGC (Fig. 2A, Table S2). Strikingly, none of the 442 AlkX homologs localize to within 20 kb of the most proximal gene cluster in that organism (Fig. 2B). In contrast, AlkZ homologs are primarily found inside or in close genomic proximity to clusters, with an average distance of roughly 2.3 kb from the nearest BGC (compared to 25 kb for AlkX). Despite their sequence similarity to AlkZ, the AlkZ2 homologs are more like AlkX in that they are also not observed within 20 kb of a BGC (Table S2).

**Figure 2.**
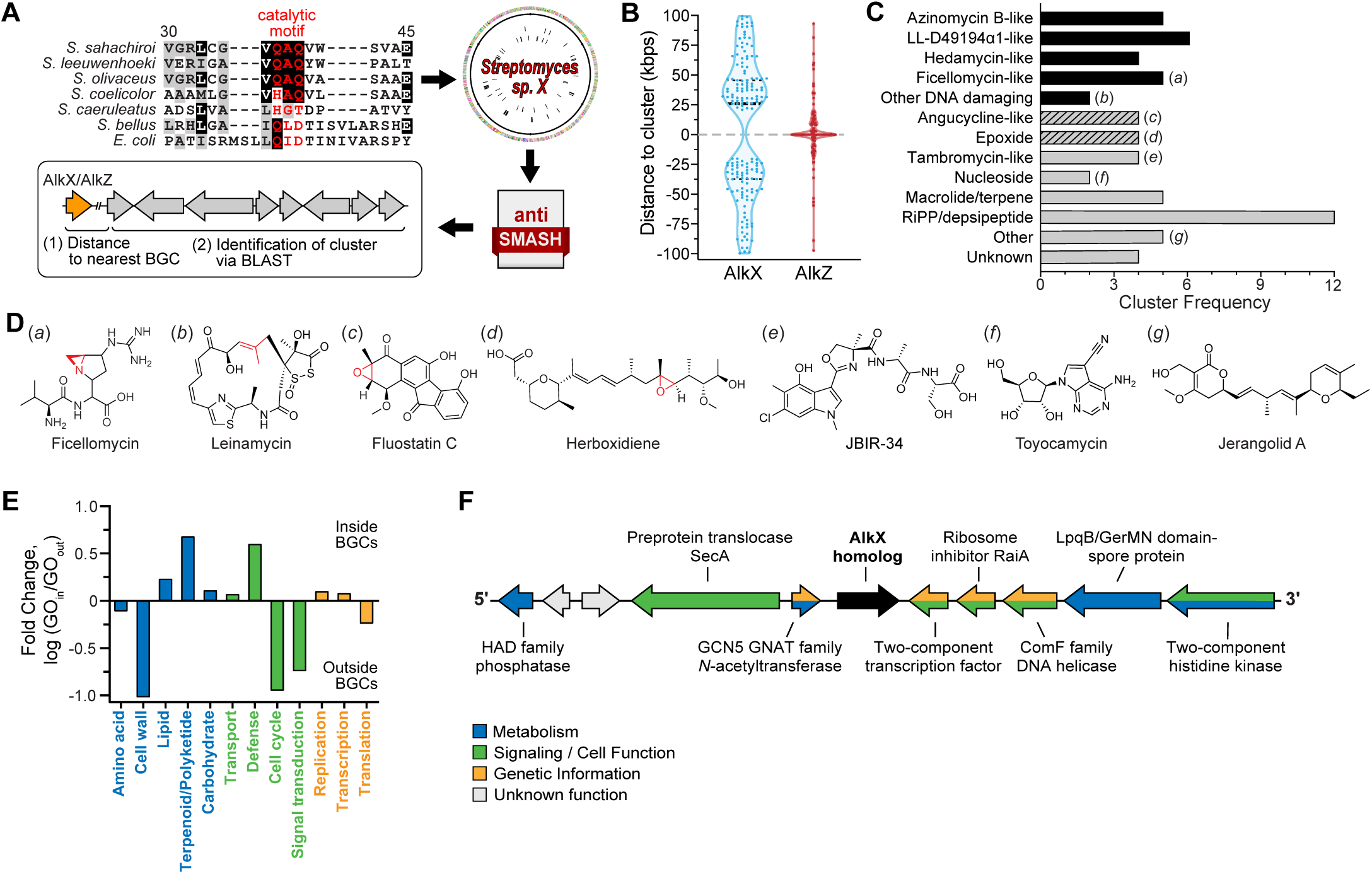
*Streptomyces* AlkZ homologs are found in diverse uncharacterized biosynthetic gene clusters. (A) Schematic depicting the work flow for identification of AlkX/AlkZ homologs in uncharacterized *Streptomyces* BGCs. Homologs were identified through the presence of the catalytic motif (red text in sequence alignment). The amino acid numbering is in relation to *S. sahachiroi* AlkZ. The corresponding *Streptomyces* genomes were input into antiSMASH, from which genomic distances between AlkX/AlkZ and the nearest BGC, as well as homologous clusters were extracted. (B) Violin plot showing the distribution of distances of AlkX (n=167) and AlkZ (n=154) homologs to the nearest BGC (in kbps; ±100 kb). The dotted line at 0 kb represents the 5’ (+) / 3’ (-) termini of the nearest BGC. Thick and thin dashed lines within the plot represent the median and upper/lower quartiles, respectively. The Chi-square significance (P) value between AlkX and AlkZ data is less than 0.0001. (C) Frequency of various types of BGCs in which AlkZ homologs were found (n=68 clusters identified). The y-axis denotes the natural product/scaffold type to which that cluster is most homologous. Black bars represent known DNA alkylators or DNA interacting metabolites, and hashed bars represent potential DNA damaging metabolites. Lowercase letters to the right of the bars correspond to structures shown in panel D. (D) Representative compounds corresponding to BGC types in panel C. Potential reactive sites are colored red. LL-D4919α1 and hedamycin structures are shown in Fig. 3. (E,F) Nearest neighbor analysis of AlkZ (E) and AlkX (F). (E) Nearest genes to AlkZ found inside and outside clusters, shown as the ratio of GO terms inside and outside, and grouped by function (blue, metabolic; green, cell signaling and function; orange, genome maintenance). (F) Representative example from *Streptomyces griseoviridis* of nearest neighbor analysis for AlkX homologs. Genes are colored according to function as in panel E (grey, unknown/hypothetical gene). These genes are invariant for all AlkX homologs, with the exception of the outermost genes, in which only one instance of variance was observed.

We found that AlkZ homologs are particularly enriched in uncharacterized *Streptomyces* BGCs, with 68 homologs localizing within a variety of different types of clusters (Fig. 2C,D; Table S3). Almost half (n=32; 47%) localize to clusters resembling those producing known DNA damaging agents, including azinomycin B (n=5), LL-D4919α1 (n=6), hedamycin (n=4), ficellomycin/vazabitide A (n=5), and C-1027/leinamycin (n=2) (Coleman et al., 2002; Dong et al., 2019; Hansen et al., 1995; Nooner et al., 2004; Reusser, 1977; Sugimoto et al., 1990). In addition, several other clusters are related to potential DNA damaging agents on the basis of a reactive epoxide functional group in the natural product, including angucycline-like molecules (n=4) herboxidiene and asukamycin. The remaining 10 uncharacterized BGCs are related to clusters that produce macrolides/terpenes, tambromycin-like compounds, and various RiPPs/depsipeptides (Fig. 2C,D).

Bacterial genes of similar function or in a particular pathway are frequently clustered into neighborhoods or operons within the genome (Mihelcic et al., 2019). Thus, we next investigated the nearest neighbors of *Streptomyces* AlkX and AlkZ genes. We collected gene ontology (GO) terms describing the biological functions of the five nearest neighbors on either side of 40 AlkX homologs, 40 AlkZ homologs inside BGCs, and 40 AlkZ homologs outside BGCs, which collectively represent ~ 15% of the total of all homologs. Biological processes were grouped into three categories—metabolism, signaling/cell function, and genetic information processing. Several key differences were found between the neighborhoods of AlkZ homologs inside versus outside clusters (Fig. 2E; Fig. S2). AlkZ genes within BGCs were more often found near terpenoid/polyketide/non-ribosomal protein synthesis and resistance/defense genes. The defense genes fell into several types: ABC transporters/permeases, α/β-fold hydrolases (VOC resistance proteins), DinB DNA-damage inducible hydrolases, and other AlkZ homologs. For those AlkZ genes found outside of BGCs, there are an abundance of neighbors involved in cell wall biosynthesis, cell cycle control, and signal transduction. In contrast, there were no significant differences between AlkZ neighbors involved in processing genetic information inside versus outside clusters. In contrast to the variation in the function of AlkZ gene neighbors, the functions of gene neighbors of AlkX homologs (outside clusters) are nearly invariant, and are composed of a variety of different gene types with no apparent functional connection between them (Fig. 2F). The functions of many of these neighbors have not been elucidated in *Streptomyces*, but some are homologous to N-acetyltransferase, a two-component transcription factor/histidine kinase, and a DNA helicase (ComF) involved in transformation competence. Thus, both the sequences and the genomic neighborhoods of AlkX homologs are relatively conserved and always found outside of BGCs, in contrast to the more variable copy number, sequence, and neighborhood of AlkZ genes prevalent within BGCs.

### Characterized BGCs containing AlkZ homologs

With the discovery that a significant proportion of AlkZ homologs reside within BGCs, we took a closer look at the nine characterized BGCs identified to contain an AlkZ homolog in the MIBiG database (Table S4). Four of these produce known DNA-alkylating agents (Fig. 3A), which contain reactive epoxide moieties like AZB that are scaffolded on diverse natural product backbones (Fig. 3A). Whereas AZB is a bifunctional alkylating agent, hedamycin (Hed), trioxacarcin A (TXN), and LL-D49194α1 (LLD) are monofunctional alkylating agents that react with N7 of guanine in specific nucleotide sequences via their epoxide rings, and also intercalate the DNA helix via their planar ring systems (Dong et al., 2019; Hansen et al., 1995; Pfoh et al., 2008). TXN and LLD clusters each contain two AlkZ paralogs (TxnU2/U4 and LldU1/U5), whereas the Hed cluster contains one (HedH4) that resides between the two polyketide synthase genes.

**Figure 3.**
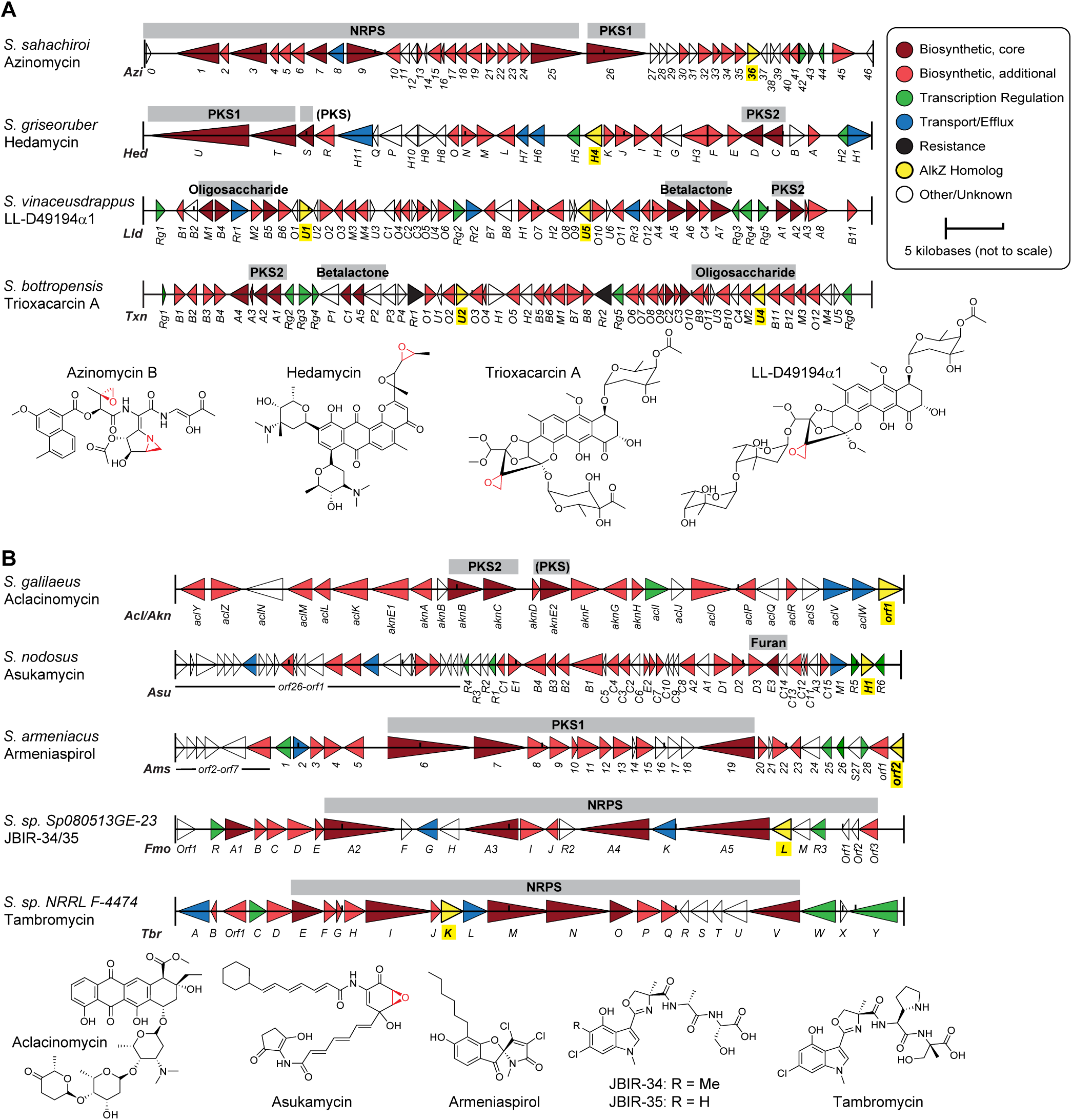
AlkZ homologs found in characterized *Streptomyces* biosynthetic gene clusters. (A,B) Gene diagrams for AlkZ-containing BGCs producing DNA alkylating agents (A) and compounds not known to alkylate DNA (B). Gene names are labeled below the cluster diagrams. The biosynthetic scaffold produced by specific genes in the cluster are labeled above the respective genes. NRPS, non-ribosomal peptide synthetase; PKS1/PKS2, type 1/2 polyketide synthase; (PKS), PKS-like. Chemical structures of the metabolites produced by each cluster are shown at the bottom of each panel.

The remaining five AlkZ-containing clusters in MIBiG produce compounds that are not known to alkylate DNA, but that share some structural characteristics with the alkylating agents described above (Fig. 3B). Aclacinomycin contains an anthracycline core surrounded by sugars that allow it to intercalate into DNA and act as a topoisomerase I poison, potentially generating downstream DNA damage (Nitiss et al., 1997). Asukamycin contains a modified PKS scaffold and an electrophilic epoxide ring and has been shown to act as both a farsenyltransferase inhibitor and a molecular glue between the UBR7 E3 ubiquitin ligase and the TP53 tumor suppressor, leading to cell death (Hara et al., 1993; Isobe et al., 2020). Armeniaspirol contains a unique chlorinated pyrrole and inhibits the AAA+ proteases ClpXP and ClpYQ leading to cell division arrest in Gram positive bacteria (Labana et al., 2021; Qiao et al., 2019). The other two BGCs produce compounds of known structure but unknown function—tambromycin and JBIR-34/35 are similar NRPS compounds containing densely substituted chlorinated indole and methyloxazoline moieties (Muliandi et al., 2014). The presence of AlkZ homologs in these clusters suggests that these compounds may be genotoxins or otherwise react with DNA, and/or that these particular AlkZ homologs may have a function outside of DNA repair.

### The AlkZ homolog within the hedamycin BGC is a DNA glycosylase that provides cellular resistance to hedamycin toxicity

The *alkZ* gene embedded within the AZB BGC provides exquisite resistance to the potent cytotoxicity of this natural product (Wang et al., 2016; Zhao et al., 2008). To determine if AlkZ homologs other than in the AZB BGC provide self-resistance to their cognate natural products, we characterized the DNA glycosylase and cellular resistance activities of HedH4 for Hed-DNA adducts. Hedamycin is a potent antibiotic/antitumor agent that induces a strong DNA damage response (Tu et al., 2004). The bisepoxide side chain alkylates the N7-position of guanines in 5’ - (C/T)G sequences (Fig. 4A), the highly oxidized aromatic polyketide intercalates the DNA helix, and two C-glycosidic linked aminosugars interact with the minor groove (Das and Khosla, 2009). We generated site-specifically labeled Hed-DNA adducts by reacting purified compound with an oligonucleotide containing a hedamycin target sequence, d(T<u>G</u>TA). The Hed-DNA adduct was stable relative to other *N*7-alkylguanine lesions as judged by thermal depurination (Fig. S3B) (Bradley et al., 2020). We assessed the ability of purified HedH4 to hydrolyze Hed-DNA adducts using a gel-based glycosylase assay that monitors alkaline cleavage of the AP site product (Bradley et al., 2020; Mullins et al., 2017). Reaction of HedH4 with Hed-DNA followed by hydroxide work-up resulted in β- and δ-elimination products consistent with production of an AP site from DNA glycosylase-mediated excision of the N-glycosidic bond of the Hed-modified nucleotide (Fig. 4A,B). We verified the identity of the excision product as Hed-guanine by HPLC/MS (Fig. 4C). To verify that the Hed-DNA product was not generated by a contaminating enzyme, we purified alanine mutants of the two glutamine residues in the QΦQ catalytic motif and tested their activity under single-turnover conditions (Fig. 4D, Fig. S3A,C). The calculated rate constant (*k*_cat_) for wild-type HedH4 was at least 7.8 ± 0.5 min^-1^ (the reaction was complete at the earliest time point). Relative to wild-type, the Q41A mutant was at least 225-fold slower (*k*_cat_ = 0.04 ± 0.01 min^-1^) and the Q43A mutant at least 10-fold slower (*k*_cat_ = 0.8 ± 0.2 min^-1^). Neither *E. coli* AlkX nor *S. sahachiroi* AlkZ displayed activity after 1 hour, indicating that Hed-DNA adducts are specifically removed by HedH4.

**Figure 4.**
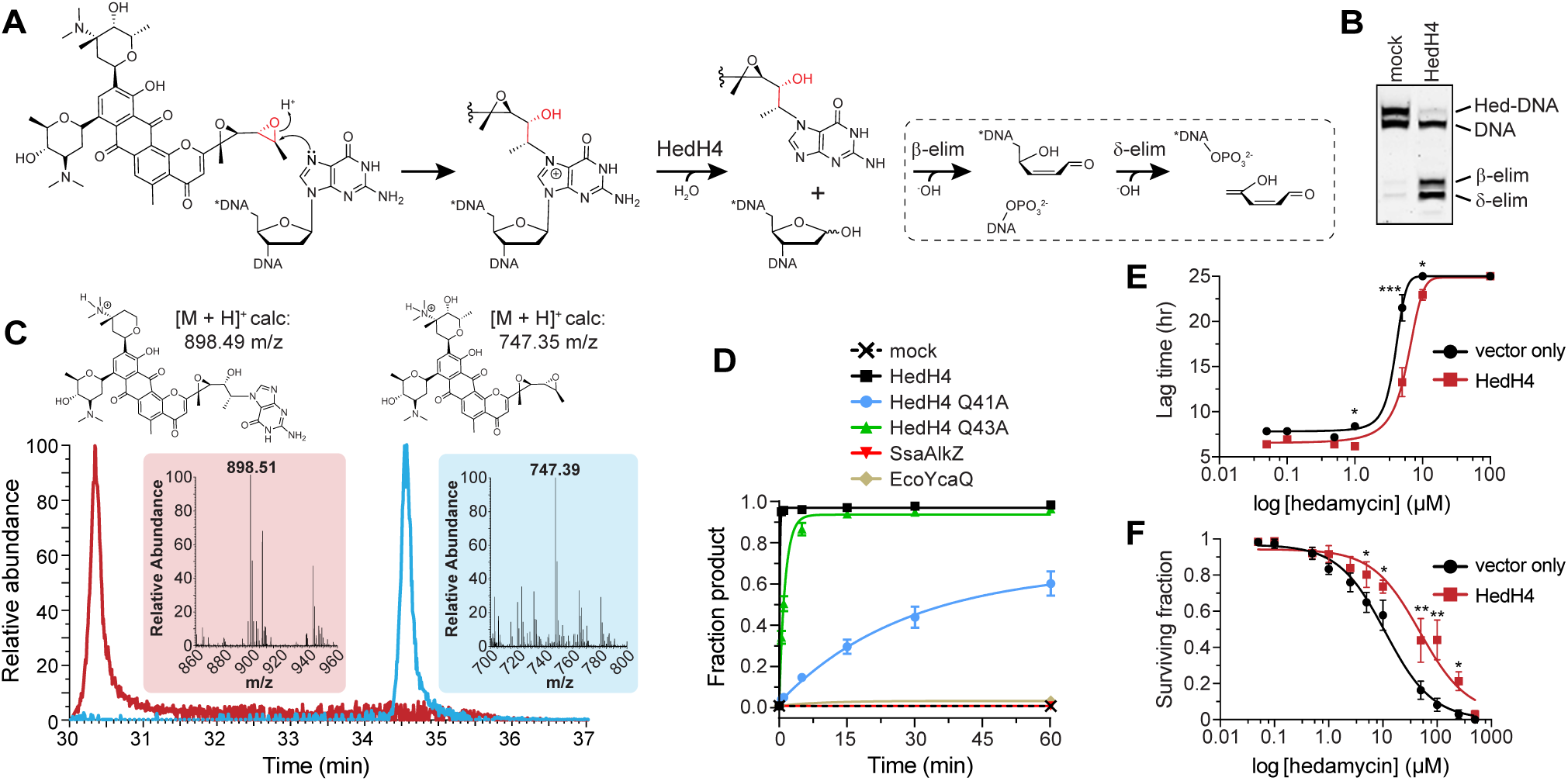
HedH4 excises hedamycin-guanine adducts from DNA and provides cellular resistance to hedamycin toxicity. (A) Hedamycin modification of deoxyguanosine in DNA forms a Hed-DNA adduct that is hydrolyzed by HedH4 to generate an abasic (AP) site in the DNA and free Hed-guanine. The reactions within the dashed line are not catalyzed by HedH4. The AP nucleotide is susceptible to base-catalyzed nicking to form shorter DNA products containing either a 3′-phospho-α,β-unsaturated aldehyde (β-elimination) or a 3′-phosphate (δ -elimination). The asterisk (*) denotes the original 5′-end of the DNA. (B) Denaturing PAGE of 5’ -Cy5-labeled Hed-DNA substrate and β- and δ -elimination products after treatment with enzyme or buffer (mock) for 1 hour, followed by NaOH to nick the AP site. The Hed-DNA reaction only goes to ~ 50% completion under our reaction conditions, as shown by the two bands of equal intensity in the mock reaction. (C) HPLC-MS analysis of hedamycin (blue) and the hedamycin-guanine excision product from reaction of HedH4 and Hed-DNA (red). Axis represents elution time (x-axis) versus relative abundance from total ion count (y-axis). Insets show mass spectra of each elution peak. (D) Wild-type and mutant HedH4 glycosylase activity for Hed-DNA. *S. sahachiroi* AlkZ and *E. coli* YcaQ (AlkX) are shown as negative controls. Data are mean ± SD (n=3). Curves were fit to a single exponential. Representative data are shown in Fig. S3C. (E) Hedamycin inhibition of *E. coli* K-12 transformed with *hedH4*/pSF-OXB1 (constitutively expressed) or empty vector pSF-OXB1. The lag time is defined as the time elapsed before cells start to grow exponentially. Data are mean ± SD (n=3). Growth curves are shown in Fig. S3E-F. Significance values were determined by unpaired t test of the mean lag time values (*: 0.05 ≤ P ≤ 0.01; ***: 0.001 ≤ P ≤ 0.0001). (E) Colony dilution assay for *E. coli* strains ±HedH4 exposed to increasing concentrations of hedamycin for 1 hour. Surviving fraction (%) is relative to untreated cells. Values are mean ± SD (n = 3). Significance values were determined by unpaired t test of the mean sensitivity values (*: 0.05 ≤ P ≤ 0.01; **: 0.01 ≤ P ≤ 0.001).

We next tested if the *hedH4* gene provides heterologous resistance to hedamycin cytotoxicity in cells. *E. coli* transformed with either vector containing *hedH4* constitutively expressed at low levels or vector alone were grown in the presence of increasing amounts of hedamycin (Fig. S3D-G). HedH4 provided a modest protection against hedamycin, as cells expressing HedH4 grew to a higher density at all hedamycin concentrations (Fig. S3F-G) and had a higher IC_50_ than cells treated with vector alone (HedH4, 5.9 μM ± 0.71; vector, 3.9 μM ± 0.36). The sensitivity differences between HedH4 and vector control were more pronounced from a colony dilution assay performed under log-phase growth conditions (Fig. 4F). Cells expressing empty vector displayed an IC_50_ value of 11.1 μM ± 1.53, while cells expressing HedH4 displayed a 4-fold reduction in sensitivity to hedamycin (48.1 μM ± 13.8). These results indicate that HedH4 is a DNA glycosylase specific for Hed-DNA adducts and provides resistance to cells exposed to the antibiotic.

## Discussion

Phylogenetic characterization of the HTH_42 superfamily within *Streptomyces* reveals three distinct subfamilies – AlkX, AlkZ, and AlkZ2. Most strikingly, *alkZ* genes, which are most prevalent in environmental microbes such as those from the phylum Actinobacteria (Fig. S1B), are highly enriched in BGCs. We found *alkZ* homologs in BGCs that produce a variety of verified and putative genotoxins, with approximately one-fifth of all *alkZ* homologs located in BGCs predicted to produce a DNA alkylating agent. We show that the AlkZ homolog, HedH4, from the hedamycin cluster excises Hed-DNA adducts and improves viability of cells grown in the presence of the hedamycin. Together with the previous example from the AZB BGC (Bradley et al., 2020; Wang et al., 2016), there is now mounting evidence that *alkZ* genes have conserved DNA repair self-resistance functions against a variety of genotoxic natural products. Indeed, we recently found that the two AlkZ homologs in both TXN and LLD clusters (TxnU2, TxnU4, LldU1, LldU5) are self-resistance glycosylases for these compounds (Chen et al.). Interestingly, AlkZ homologs were also found in BGCs that by homology were not expected to produce DNA alkylators or other genotoxins. These clusters could have additional uncharacterized enzymes such as cytochrome P450s, sulfate adenyltransferases, or epoxidases that could activate the natural products into DNA alkylators (Thibodeaux et al., 2012). Alternatively, AlkZ proteins in these clusters could have regulatory or protective roles outside of DNA repair. Consistent with their role in resistance, the *alkZ* genes found inside BGCs frequently localize around a variety of other resistance genes. Moreover, the relatively high copy number and low sequence conservation of AlkZ proteins is consistent with increased expression or possible horizontal gene transfer events that enable these enzymes to evolve specificity for particular natural product (Hastings et al., 2009).

In contrast to the genotoxin-specific *alkZ* genes, *alkX* and *alkZ2* are always found outside clusters and thus are likely to provide a more general role in protecting the genome against environmental genotoxins, similar to that shown for *E. coli* AlkX (YcaQ) (Bradley et al., 2020). AlkX homologs and their gene neighborhoods very highly conserved, suggesting they play a critical role as part of a unified pathway (Rogozin et al., 2002). Although the particular pathway is unclear, the presence of a two-component transcription factor/kinase and ComF DNA helicase within the AlkX neighborhood in *Streptomyces* also hints at a signaling network for DNA uptake (Londono-Vallejo and Dubnau, 1993; Turgay et al., 1998; Veening and Blokesch, 2017). Similarly, *E. coli alkX* is localized in a four-gene operon involved in cell wall biosynthesis and transformation competence (Bradley et al., 2020). Continued exploration of the gene neighborhoods of *alkX* and *alkZ* beyond *Streptomyces* will reveal a deeper understanding of the cellular roles played by these enzymes. This will be especially important for AlkX, which are prevalent in human pathogens or commensal microbes (Wang et al., 2016).

Resistance genome mining has emerged as a critical bioinformatically driven pipeline to discover novel natural products and gene clusters in several organisms (Panter et al., 2018; Yan et al., 2018). A key benefit of resistance genome mining is the dramatically decreased candidate pool as a result of targeted identification of gene clusters containing a resistance gene. Generally, these methods require a basic understanding of the resistance mechanisms involved. We sought to use this approach for the first time to hunt for BGCs that produce alkylating genotoxins, using prior knowledge of the DNA repair functions of *S. sahachiroi* AlkZ within the AZB cluster (Bradley et al., 2020; Mullins et al., 2017; Wang et al., 2016). In this study, we examined 435 *Streptomyces* species for BGCs within which AlkZ was located and found 62 uncharacterized clusters that are candidates for targeted elucidation of their products. Characterization of these orphan clusters could provide new analogs or types of DNA alkylating/damaging secondary metabolites, an important step in developing new antitumor or antibiotic treatments. This classification of AlkX/AlkZ proteins in *Streptomyces* is an important first step in understanding their evolutionary connection to each other and to BGCs of different types, and demonstrates that targeted resistance genome mining is a viable approach to discover novel genotoxins and resistance mechanisms from uncharacterized BGCs.

## Materials and Methods

### Reagents

The pBG102 expression vector was obtained from the Vanderbilt University Center for Structural Biology, and pSF-OXB1 was purchased from Millipore Sigma. DNA oligonucleotides (Table S5) were purchased from Integrated DNA Technologies. *Escherichia coli* K-12 wild-type strain was purchased from the Keio *E. coli* knockout collection (Dharmacon, GE Healthcare). Hedamycin was obtained from the National Cancer Institute’s Developmental Therapeutic Program (NCI DTP) Open Compound Repository (NSC 70929). Unless otherwise noted, all other chemicals were purchased from Sigma-Aldrich.

### Taxonomy and phylogeny of Streptomyces AlkX and AlkZ

To identify homologs of *alkX and alkZ* in *Streptomyces*, the protein sequences for AlkX (GenBank accession number QHB65847.1) and AlkZ (GenBank accession number ABY83174.1) were used for tBLASTn and BLASTp searches (BLAST+ v2.11.0) against all *Streptomyces* genomes (taxid:1883). Searches were run with the BLOSUM62 matrix, 1000 maximum target sequences, and 0.05 threshold using an e-value and identity cutoff of 10^−4^ and 25%, respectively. All hits were verified for the presence of the (H/Q)Φ (D/Q) catalytic motif, during which the (H/Q)Φ (S/T)(D/E) (AlkZ2) variant was identified. Truncated genes, poor sequence quality genes, and pseudogenes were eliminated. Additional sequences were obtained by searching the Pfam database v33.1 (El-Gebali et al., 2019) for *Streptomyces* HTH_42 superfamily members (PF06224). Sequences from Pfam were sorted according to their domain classes (Fig. S1A), and only sequences from Class 1 with >75% coverage were included. Protein sequences were aligned using EMBL-EBI Clustal OmegaW or MAFFT v7 using default parameters (Katoh et al., 2019; Madeira et al., 2019). The evolutionary history of AlkX/AlkZ sequences were reconstructed using IQTREE2 with default settings (Minh et al., 2020), and the phylogenetic tree was assembled with the Interactive Tree of Life (v5) phylogeny display tool (Letunic and Bork, 2019). Sequence logos were generated with WebLogo v2.8.2 (Crooks et al., 2004). The copy number frequency and coincidence of AlkX/AlkZ in the same genome was determined by manually counting the number and identity of homologs in each species. A list of all *alkX*/*alkZ/alkZ2* homologs and *Streptomyces* genomes analyzed in this study can be found in Table S1.

### Identification of AlkZ homologs in known biosynthetic gene clusters

To find AlkZ homologs in verified and/or published BGCs, we searched MIBiG v2.0 for the azinomycin B BGC (BGC0000960) from *S. sahachiroi* (Kautsar et al., 2020; Zhao et al., 2008), followed by an iterative search using the *MIBiG Hits* function until no more hits were obtained. The homologs TxnU2 and TxnU4 were identified from the initial BLAST search within the deposited NCBI trioxacarcin BGC sequence (Zhang et al., 2015). The homolog within the aclacinomycin BGC was also identified in the initial BLAST search as appearing in proximity to aclacinomycin biosynthesis genes. Closer inspection of the published sequence for the aclacinomycin BGC (GenBank accession number AB008466.1) revealed an AlkZ homolog (Orf1) located immediately 3’ of the cluster (Chung et al., 2002). A detailed list of the AlkZ homologs in known BGCs can be found in Table S3.

### Identification of AlkZ homologs in uncharacterized biosynthetic gene clusters

To determine the physical distance in base pairs between the genomic coordinates of *alkZ* homologs and those of BGCs present in the genome assemblies of *Streptomyces* (average number of scaffolds: 96.30; minimum: 1; maximum: 1,956), we first predicted the BGCs in each genome using antiSMASH v5.1.0 (Blin et al., 2019) with the *taxon* parameter set to *bacteria*. Using the BGC sequences identified from antiSMASH and *alkZ* homolog sequences, a custom python script using Biopython (Cock et al., 2009) determined the shortest base pair distance between the physical location of the *alkX/alkZ* homolog and the location of the nearest BGC on the same scaffold (less than 2 Mbps away). To be considered within a BGC, the homolog had to be observed within 5 genes or 2 kb of the nearest cluster. Known Cluster BLAST was performed within antiSMASH to determine the most similar BGC to the unknown clusters, and the result with the highest percentage of similar genes was recorded as the most similar cluster. A detailed list of the genome information, cluster IDs, and the closest 3’ and/or 5’ BGC can be found in Table S5.

### Gene ontology analysis

To identify GO terms for nearest neighbors identified through BLAST, Pfam, and MIBiG searches, we randomly chose 40 homologs each of AlkZ inside BGCs, AlkZ outside BGCs, and AlkX, which represent ~ 10% of the sequences for each. Amino acid sequences for the five genes on both sides of the *alkX*/*alkZ* genes were downloaded from the NCBI database, for a total of 400 neighbors for each of the three classes. Cellular functions of any already annotated genes in the NCBI database were identified and recorded. The downloaded sequences were then run through the GhostKOALA (v2.2) and eggNOG (v5.0) GO annotation databases (Huerta-Cepas et al., 2019; Kanehisa et al., 2016). After known GO terms for all gene neighbors were identified, proteins were categorized by biological processes and molecular functions, and the values for these terms were used to create the GO term distributions. Proteins that had multiple GO terms associated with them were counted into each class of terms. A list of all proteins and their annotated GO terms can be found in Tables S6-S11.

### Protein purification

AlkX and AlkZ were purified as previously described (Bradley et al., 2020). *Streptomyces caeruleatus alkZ2* and *Streptomyces griseoruber hedH4* genes were synthesized by GenScript and cloned into pBG102. The N-terminal His_6_-SUMO fusion protein was overexpressed in *Escherichia coli* Tuner (DE3) cells at 16°C for 18 h in LB medium supplemented with 30 μg/mL kanamycin and 50 μM isopropyl β-D-1-thiogalactopyranoside (IPTG). Cells were lysed by sonication and cell debris removed by centrifugation at 45,000 × g at 4°C for 30 m. Clarified lysate was passed over Ni-NTA agarose equilibrated in buffer A (50 mM Tris• HCl pH 8.5, 500 mM NaCl, 25 mM imidazole, and 10% (vol/vol) glycerol) and protein eluted in 250 mM imidazole/buffer A. Protein fractions were pooled and supplemented with 0.1 mM EDTA and 1 mM tris(2-carboxyethyl)phosphine (TCEP) before incubation with 0.5 mg Rhinovirus 3C protease (PreScission) at 4°C overnight. Cleaved protein was diluted 10-fold in buffer B (50 mM Tris• HCl pH 8.5, 10% (vol/vol) glycerol, 0.1 mM TCEP, and 0.1 mM EDTA) and purified by heparin sepharose using a 0–1 M NaCl/buffer B linear gradient. Fractions were pooled and passed over Ni-NTA agarose in buffer A, concentrated and filtered, and buffer exchanged into buffer C (20 mM Tris• HCl pH 8.5, 100 mM NaCl, 5% (vol/vol) glycerol, 0.1 mM TCEP, and 0.1 mM EDTA). Protein was concentrated to 4 mg/mL, flash-frozen in liquid nitrogen, and stored at −80°C. Mutant protein expression vectors were generated using the Q5 Mutagenesis Kit (New England BioLabs) and proteins overexpressed and purified the same as wild-type.

### DNA glycosylase activity

DNA substrates containing a single N7-methyl-2′-deoxyguanosine lesion and a 5’ -Cy5 fluorophore were prepared as described previously (Mullins et al., 2013). DNA substrates containing a single hedamycin-guanosine adduct were prepared by annealing DNA with the sequence 5’ -Cy5-d(AATATTAATAAT<u>G</u>TAATTTAAATTA) to the complementary unlabeled oligo. Hedamycin was dissolved in DMSO to a concentration of 5 mM. 100 μM DNA was incubated with 200 μM hedamycin in 10% methanol and 20% DMSO at 4°C on ice in the dark for 24 h. Unreacted drug was removed using an Illustra G-25 spin column (GE Healthcare) equilibrated in TE buffer (10 mM Tris• HCl pH 8.0, 1 mM EDTA pH 8.0), and the DNA was stored at −80°C.

In each glycosylase reaction, 1 μM enzyme was incubated with 50 nM DNA in glycosylase buffer (50 mM HEPES pH 8.5, 100 mM KCl, 1 mM EDTA, and 10% (vol/vol) glycerol) at 25°C. At various time points, 4-μL aliquots were added to 1 μL of 1M NaOH and heated at 70°C for 2 m. Samples were denatured at 70°C for 5 m in 5 mM EDTA pH 8.0, 80% (wt/vol) formamide, and 1 mg/mL blue dextran prior to electrophoresis on a 20% (wt/vol) acrylamide/8 M urea sequencing gel at 40 watts for 1 hour in 0.5 × TBE buffer (45 mM Tris, 45 mM borate, and 1 mM EDTA pH 8.0). Gels were imaged on a Typhoon Trio variable mode imager (GE Healthcare) using 633-nm excitation/670-nm emission fluorescence for Cy5, and bands were quantified with ImageQuant (GE Healthcare). All excision assays were performed in triplicate.

### HPLC-MS analysis of hedamycin and hedamycin-guanine

HPLC was performed on an Agilent Series 1100 system equipped with an analytical SymmetryShield RP-C18 column (3.5 μm, 4.6 mm × 7.5 mm, 100 Å pore size) and using a linear gradient from 90% buffer A (10 mM ammonium formate) / 10% buffer B (100% methanol) to 100% B over 40 m and a flow rate of 0.4 mL/m. Hedamycin was diluted to 50 μM in 10% methanol and stored on ice prior to HPLC injection. To analyze the product of HedH4 activity, Hed-DNA was diluted to 10 μM in glycosylase buffer and reacted with 50 μM HedH4 for 1 h at room temperature before injection. Mass spectrometry was performed with an LTQ Orbitrap XL Hybrid FT Mass Spectrometer (Thermo Fisher Scientific) in positive ion mode from 300-1000 m/z.

### Cellular assays for hedamycin resistance

The *hedH4* wild-type gene was sub-cloned from pBG102 into pSF-OXB1 using NcoI and XbaI restriction sites. The pSF-OXB1 vector contains a kanamycin resistance gene and allows for constitutive low-level expression from a modified AraBAD promoter. pSF-OXB1 and HedH4/pSF-OXB1 were transformed into *E. coli* K-12 cells. Cloning of *hedH4* was confirmed by sequencing, restriction digest using NcoI-HF/XbaI (Fig. S4C), and colony PCR of K-12 transformants using the HedH4 NcoI and XbaI primers (Fig. S4D, Table S1). Cultures were grown at 37°C in LB media supplemented with 30 μg/mL Kan. Growth curves were generated by diluting overnight cultures to 0.01 OD_600_ in LB/Kan supplemented with 0 nM-100 μM hedamycin in a 96-well flat-bottom plate. The plate was incubated at 30°C with shaking for 24 h and cell density was measured at 600 nm every 20 min using a Bio-Tek Synergy 2 microplate reader. IC_50_ values were determined from a fit to the equation, Lag time = Min_lag_ + (Max_lag_-Min_lag_)/(1+(IC_50_/[Hed])^*h*^). Growths were performed in triplicate.

*E. coli* survival curves after hedamycin treatment were performed using a colony dilution assay. A saturated overnight LB/Kan culture from a single colony was diluted to 0.01 OD_600_ in 1 mL fresh LB/Kan media and grown to 0.6 OD_600_ at 37°C. The cells were treated with various concentrations of hedamycin for 1 h at 37°C. Treated cells were transferred to fresh LB/Kan media, serially diluted by 10^−6^ in LB/Kan media, and 100 μL of diluted cells were plated on LB/Kan agar plates and grown at 37°C overnight. Colonies were counted the next morning and the CFU/mL culture was determined. The percent survival was calculated as CFU/mL(treated) / CFU/mL(untreated). Curves were plotted on a logarithmic scale and IC_50_ values determined by non-linear regression fits to the data. Growths were performed in triplicate.

## Abbreviations

AZB: azinomycin B
Hed: hedamycin
BER: base excision repair
BGC: biosynthetic gene cluster
GO: gene ontology
ICL: interstrand crosslink
RGM: resistance genome mining

## Acknowledgements

This work was supported by grants from the National Institutes of Health (R01GM131071) and the National Science Foundation (MCB-1928918) to B.F.E. Research in A.R.’s lab is supported by grants from the National Science Foundation (DEB-1442113 and DEB-2110404), the National Institutes of Health/National Institute of Allergy and Infectious Diseases (R56 AI146096), and the Burroughs Wellcome Fund. N.P.B. was supported by the Vanderbilt Training Program in Environmental Toxicology (NIH T32ES007028) and an NSF Graduate Research Fellowship (DGE-1445197). J.L.S. and A.R. are supported by the Howard Hughes Medical Institute through the James H. Gilliam Fellowships for Advanced Study program.

## Author Contributions

B.F.E. and N.P.B. conceived of the study; N.P.B., K.L.W., and J.L.S. performed experiments; N.P.B., K.L.W., and B.F.E. designed experiments, analyzed data, and wrote the manuscript. All authors provided feedback on the interpretation of the results and the manuscript.

## References

Belknap, K.C., Park, C.J., Barth, B.M., and Andam, C.P. (2020). Genome mining of biosynthetic and chemotherapeutic gene clusters in Streptomyces bacteria. Sci Rep 10, 2003.

Bizanek, R., McGuinness, B.F., Nakanishi, K., and Tomasz, M. (1992). Isolation and Structure of an Intrastrand Cross-Link Adduct of Mitomycin C and DNA. Biochemistry 31, 3084–3091.

Blin, K., Shaw, S., Steinke, K., Villebro, R., Ziemert, N., Lee, S.Y., Medema, M.H., and Weber, T. (2019). antiSMASH 5.0: updates to the secondary metabolite genome mining pipeline. Nucleic Acids Res 47, W81–W87.

Boger, D.L., and Garbaccio, R.M. (1997). Catalysis of the CC-1065 and Duocarmycin DNA Alkylation Reaction: DNA Binding Induced Conformational Change in the Agent Results in Activation Bioorganic and Medicinal Chemistry 5, 263–276.

Bradley, N.P., Washburn, L.A., Christov, P.P., Watanabe, C.M.H., and Eichman, B.F. (2020). Escherichia coli YcaQ is a DNA glycosylase that unhooks DNA interstrand crosslinks. Nucleic Acids Res 48, 7005–7017.

Chen, X., Bradley, N.P., Lub, W., Wahl, K.L., Zhang, M., Yuan, H., Hou, X.F., Eichman, B.F., and Tang, G.L. Base excision repair system targeting DNA adducts of antibiotics trioxacarcin/LL-D49194 for self-resistance. Manuscript in preparation.

Chumduri, C., Gurumurthy, R.K., Zietlow, R., and Meyer, T.F. (2016). Subversion of host genome integrity by bacterial pathogens. Nat Rev Mol Cell Biol 17, 659–673.

Chung, J.Y., Fujii, I., Harada, S., Sankawa, U., and Ebizuka, Y. (2002). Expression, purification, and characterization of AknX anthrone oxygenase, which is involved in aklavinone biosynthesis in Streptomyces galilaeus. J Bacteriol 184, 6115–6122.

Cock, P.J., Antao, T., Chang, J.T., Chapman, B.A., Cox, C.J., Dalke, A., Friedberg, I., Hamelryck, T., Kauff, F., Wilczynski, B., et al. (2009). Biopython: freely available Python tools for computational molecular biology and bioinformatics. Bioinformatics 25, 1422–1423.

Coleman, R.S., Perez, R.J., Burk, C.B., and Navarro, A. (2002). Studies on the Mechanism of Action of Azinomycin B: Definition of Regioselectivity and Sequence Selectivity of DNA Cross-Link Formation and Clarification of the Role of the Naphthoate. Journal of the American Chemical Society 124, 13008–13017.

Crooks, G.E., Hon, G., Chandonia, J.M., and Brenner, S.E. (2004). WebLogo: a sequence logo generator. Genome Res 14, 1188–1190.

Cundliffe, E., and Demain, A.L. (2010). Avoidance of suicide in antibiotic-producing microbes. J Ind Microbiol Biotechnol 37, 643–672.

D’Costa, V.M., McGrann, K.M., Hughes, D.W., and Wright, G.D. (2006). Sampling the antibiotic resistome. Science 311, 374–377.

Das, A., and Khosla, C. (2009). In vivo and in vitro analysis of the hedamycin polyketide synthase. Chem Biol 16, 1197–1207.

Demain, A.L., and Sanchez, S. (2009). Microbial drug discovery: 80 years of progress. J Antibiot (Tokyo) 62, 5–16.

Dong, L., Shen, Y., Hou, X.F., Li, W.J., and Tang, G.L. (2019). Discovery of Druggability-Improved Analogues by Investigation of the LL-D49194α1 Biosynthetic Pathway. Org Lett 21, 2322–2325.

El-Gebali, S., Mistry, J., Bateman, A., Eddy, S.R., Luciani, A., Potter, S.C., Qureshi, M., Richardson, L.J., Salazar, G.A., Smart, A., et al. (2019). The Pfam protein families database in 2019. Nucleic Acids Res 47, D427–D432.

Furuya, K., and Hutchinson, C.R. (1998). The DrrC protein of Streptomyces peucetius, a UvrA-like protein, is a DNA-binding protein whose gene is induced by daunorubicin. FEMS Microbiol Lett 168, 243–249.

Galm, U., Hager, M.H., Van Lanen, S.G., Ju, J., Thorson, J.S., and Shen, B. (2005). Antitumor antibiotics: bleomycin, enediynes, and mitomycin. Chem Rev 105, 739–758.

Gates, K.S. (2009). An Overview of Chemical Processes That Damage Cellular DNA: Spontaneous Hydrolysis, Alkylation, and Reactions with Radicals. Chemical Research in Toxicology 22, 1747–1760.

Hansen, M., and Hurley, L. (1995). Altromycin B Threads the DNA Helix Interacting with Both the Major and the Minor Grooves To Position Itself for Site-Directed Alkylation of Guanine N7. J Am Chem Soc 117, 2421–2429.

Hansen, M., Yun, S., and Hurley, L. (1995). Hedamycin intercalates the DNA helix and, through carbohydrate-mediated recognition in the minor groove, directs N-alkylation of guanine in the major groove in a sequence-specific manner Chemistry & Biology 2, 229–240.

Hara, M., Akasaka, K., Akinaga, S., Okabe, M., Nakano, H., Gomez, R., Wood, D., Uh, M., and Tamanoi, F. (1993). Identification of Ras farnesyltransferase inhibitors by microbial screening. Proc Natl Acad Sci U S A 90, 2281–2285.

Hastings, P.J., Lupski, J.R., Rosenberg, S.M., and Ira, G. (2009). Mechanisms of change in gene copy number. Nat Rev Genet 10, 551–564.

Huang, M., Lu, J.J., and Ding, J. (2021). Natural Products in Cancer Therapy: Past, Present and Future. Nat Prod Bioprospect 11, 5–13.

Huerta-Cepas, J., Szklarczyk, D., Heller, D., Hernandez-Plaza, A., Forslund, S.K., Cook, H., Mende, D.R., Letunic, I., Rattei, T., Jensen, L.J., et al. (2019). eggNOG 5.0: a hierarchical, functionally and phylogenetically annotated orthology resource based on 5090 organisms and 2502 viruses. Nucleic Acids Res 47, D309–D314.

Isobe, Y., Okumura, M., McGregor, L.M., Brittain, S.M., Jones, M.D., Liang, X., White, R., Forrester, W., McKenna, J.M., Tallarico, J.A., et al. (2020). Manumycin polyketides act as molecular glues between UBR7 and P53. Nat Chem Biol 16, 1189–1198.

Jacob, C., and Weissman, K.J. (2017). Unpackaging the Roles of Streptomyces Natural Products. Cell Chem Biol 24, 1194–1195.

Kanehisa, M., Sato, Y., and Morishima, K. (2016). BlastKOALA and GhostKOALA: KEGG Tools for Functional Characterization of Genome and Metagenome Sequences. J Mol Biol 428, 726–731.

Katoh, K., Rozewicki, J., and Yamada, K.D. (2019). MAFFT online service: multiple sequence alignment, interactive sequence choice and visualization. Brief Bioinform 20, 1160–1166.

Kautsar, S.A., Blin, K., Shaw, S., Navarro-Munoz, J.C., Terlouw, B.R., van der Hooft, J.J.J., van Santen, J.A., Tracanna, V., Suarez Duran, H.G., Pascal Andreu, V., et al. (2020). MIBiG 2.0: a repository for biosynthetic gene clusters of known function. Nucleic Acids Res 48, D454–D458.

Kersten, R.D., and Weng, J.K. (2018). Gene-guided discovery and engineering of branched cyclic peptides in plants. Proc Natl Acad Sci U S A 115, E10961–E10969.

Kjaerbolling, I., Vesth, T., and Andersen, M.R. (2019). Resistance Gene-Directed Genome Mining of 50 Aspergillus Species. mSystems 4.

Kwok, Y., Zeng, Q., and Hurley, L. (1998). Topoisomerase II-mediated site-directed alkylation of DNA by psorospermin and its use in mapping other topoisomerase II poison binding sites. Proc Natl Acad Sci U S A 95, 13531–13536.

Labana, P., Dornan, M.H., Lafreniere, M., Czarny, T.L., Brown, E.D., Pezacki, J.P., and Boddy, C.N. (2021). Armeniaspirols inhibit the AAA+ proteases ClpXP and ClpYQ leading to cell division arrest in Gram-positive bacteria. Cell Chem Biol.

Law, J.W., Law, L.N., Letchumanan, V., Tan, L.T., Wong, S.H., Chan, K.G., Ab Mutalib, N.S., and Lee, L.H. (2020). Anticancer Drug Discovery from Microbial Sources: The Unique Mangrove Streptomycetes. Molecules 25.

Letunic, I., and Bork, P. (2019). Interactive Tree Of Life (iTOL) v4: recent updates and new developments. Nucleic Acids Res 47, W256–W259.

Lomovskaya, N., Hong, S.K., Kim, S.U., Fonstein, L., Furuya, K., and Hutchinson, R.C. (1996). The Streptomyces peucetius drrC gene encodes a UvrA-like protein involved in daunorubicin resistance and production. J Bacteriol 178, 3238–3245.

Londono-Vallejo, J.A., and Dubnau, D. (1993). comF, a Bacillus subtilis late competence locus, encodes a protein similar to ATP-dependent RNA/DNA helicases. Mol Microbiol 9, 119–131.

Ma, L., Sun, S., Yuan, Z., Deng, Z., Tang, Y., and Yu, Y. (2020). Three putative DNA replication/repair elements encoding genes confer self-resistance to distamycin in Streptomyces netropsis. Acta Biochim Biophys Sin (Shanghai) 52, 91–96.

Madeira, F., Park, Y.M., Lee, J., Buso, N., Gur, T., Madhusoodanan, N., Basutkar, P., Tivey, A.R.N., Potter, S.C., Finn, R.D., et al. (2019). The EMBL-EBI search and sequence analysis tools APIs in 2019. Nucleic Acids Res 47, W636–W641.

Mihelcic, M., Smuc, T., and Supek, F. (2019). Patterns of diverse gene functions in genomic neighborhoods predict gene function and phenotype. Sci Rep 9, 19537.

Minh, B.Q., Schmidt, H.A., Chernomor, O., Schrempf, D., Woodhams, M.D., von Haeseler, A., and Lanfear, R. (2020). IQ-TREE 2: New Models and Efficient Methods for Phylogenetic Inference in the Genomic Era. Mol Biol Evol 37, 1530–1534.

Muliandi, A., Katsuyama, Y., Sone, K., Izumikawa, M., Moriya, T., Hashimoto, J., Kozone, I., Takagi, M., Shin-ya, K., and Ohnishi, Y. (2014). Biosynthesis of the 4-methyloxazoline-containing nonribosomal peptides, JBIR-34 and -35, in Streptomyces sp. Sp080513GE-23. Chem Biol 21, 923–934.

Mullins, E.A., Rodriguez, A.A., Bradley, N.P., and Eichman, B.F. (2019). Emerging Roles of DNA Glycosylases and the Base Excision Repair Pathway. Trends Biochem Sci 44, 765–781.

Mullins, E.A., Rubinson, E.H., Pereira, K.N., Calcutt, M.W., Christov, P.P., and Eichman, B.F. (2013). An HPLC-tandem mass spectrometry method for simultaneous detection of alkylated base excision repair products. Methods 64, 59–66.

Mullins, E.A., Warren, G.M., Bradley, N.P., and Eichman, B.F. (2017). Structure of a DNA glycosylase that unhooks interstrand cross-links. Proc Natl Acad Sci U S A 114, 4400–4405.

Mungan, M.D., Alanjary, M., Blin, K., Weber, T., Medema, M.H., and Ziemert, N. (2020). ARTS 2.0: feature updates and expansion of the Antibiotic Resistant Target Seeker for comparative genome mining. Nucleic Acids Res 48, W546–W552.

Ng, T.L., Rohac, R., Mitchell, A.J., Boal, A.K., and Balskus, E.P. (2019). An N-nitrosating metalloenzyme constructs the pharmacophore of streptozotocin. Nature 566, 94–99.

Nitiss, J.L., Pourquier, P., and Pommier, Y. (1997). Aclacinomycin A Stabilizes Topoisomerase I Covalent Complexes. Cancer Research 57, 4564–4569.

Nooner, T., Dutta, S., and Gates, K.S. (2004). Chemical Properties of the Leinamycin-Guanine Adduct in DNA. Chemical Research in Toxicology 17, 942–949.

Panter, F., Krug, D., Baumann, S., and Muller, R. (2018). Self-resistance guided genome mining uncovers new topoisomerase inhibitors from myxobacteria. Chem Sci 9, 4898–4908.

Parrish, J.P., Kastrinsky, D.B., Wolkenberg, S.E., Igarashi, Y., and Boger, D.L. (2003). DNA Alkylation Properties of Yatakemycin. Journal of the American Chemical Society 125, 10971–10976.

Pfoh, R., Laatsch, H., and Sheldrick, G.M. (2008). Crystal structure of trioxacarcin A covalently bound to DNA. Nucleic Acids Res 36, 3508–3514.

Prija, F., Srinivasan, P., Das, S., Kattusamy, K., and Prasad, R. (2017). DnrI of Streptomyces peucetius binds to the resistance genes, drrAB and drrC but is activated by daunorubicin. J Basic Microbiol 57, 862–872.

Procopio, R.E., Silva, I.R., Martins, M.K., Azevedo, J.L., and Araujo, J.M. (2012). Antibiotics produced by Streptomyces. Braz J Infect Dis 16, 466–471.

Qiao, Y., Yan, J., Jia, J., Xue, J., Qu, X., Hu, Y., Deng, Z., Bi, H., and Zhu, D. (2019). Characterization of the Biosynthetic Gene Cluster for the Antibiotic Armeniaspirols in Streptomyces armeniacus. J Nat Prod 82, 318–323.

Reusser, F. (1977). Ficellomycin and feldamycin; inhibitors of bacterial semiconservative DNA replication. Biochemistry 16, 3406–3412.

Rogozin, I.A., Makarova, K.S., Murvai, J., Czabarka, E., Wolf, W.S., Tatusov, R.L., Szekely, L.A., and Koonin, E.V. (2002). Connected gene neighborhoods in prokaryotic genomes. Nucleic Acids Res 30, 2212–2223.

Seyedsayamdost, M.R. (2019). Toward a global picture of bacterial secondary metabolism. J Ind Microbiol Biotechnol 46, 301–311.

Shigdel, U.K., Lee, S.J., Sowa, M.E., Bowman, B.R., Robison, K., Zhou, M., Pua, K.H., Stiles, D.T., Blodgett, J.A.V., Udwary, D.W., et al. (2020). Genomic discovery of an evolutionarily programmed modality for small-molecule targeting of an intractable protein surface. Proc Natl Acad Sci U S A 117, 17195–17203.

Skinnider, M.A., Dejong, C.A., Rees, P.N., Johnston, C.W., Li, H., Webster, A.L., Wyatt, M.A., and Magarvey, N.A. (2015). Genomes to natural products PRediction Informatics for Secondary Metabolomes (PRISM). Nucleic Acids Res 43, 9645–9662.

Sugimoto, Y., Otani, T., Oie, S., Wierzba, K., and Yamada, Y. (1990). Mechanism of action of a new macromolecular antitumor antibiotic, C-1027. J Antibiot (Tokyo) 43, 417–421.

Tenconi, E., and Rigali, S. (2018). Self-resistance mechanisms to DNA-damaging antitumor antibiotics in actinobacteria. Curr Opin Microbiol 45, 100–108.

Thaker, M.N., Wang, W., Spanogiannopoulos, P., Waglechner, N., King, A.M., Medina, R., and Wright, G.D. (2013). Identifying producers of antibacterial compounds by screening for antibiotic resistance. Nat Biotechnol 31, 922–927.

Thibodeaux, C.J., Chang, W.C., and Liu, H.W. (2012). Enzymatic chemistry of cyclopropane, epoxide, and aziridine biosynthesis. Chem Rev 112, 1681–1709.

Tu, L.C., Melendy, T., and Beerman, T.A. (2004). DNA damage responses triggered by a highly cytotoxic monofunctional DNA alkylator, hedamycin, a pluramycin antitumor antibiotic. Molecular Cancer Therapeutics 3, 577–585.

Turgay, K., Hahn, J., Burghoorn, J., and Dubnau, D. (1998). Competence in Bacillus subtilis is controlled by regulated proteolysis of a transcription factor. EMBO J 17, 6730–6738.

Tyc, O., Song, C., Dickschat, J.S., Vos, M., and Garbeva, P. (2017). The Ecological Role of Volatile and Soluble Secondary Metabolites Produced by Soil Bacteria. Trends Microbiol 25, 280–292.

Veening, J.W., and Blokesch, M. (2017). Interbacterial predation as a strategy for DNA acquisition in naturally competent bacteria. Nat Rev Microbiol 15, 629.

Wang, S., Liu, K., Xiao, L., Yang, L., Li, H., Zhang, F., Lei, L., Li, S., Feng, X., Li, A., et al. (2016). Characterization of a novel DNA glycosylase from S. sahachiroi involved in the reduction and repair of azinomycin B induced DNA damage. Nucleic Acids Res 44, 187–197.

Xu, H., Huang, W., He, Q.L., Zhao, Z.X., Zhang, F., Wang, R., Kang, J., and Tang, G.L. (2012). Self-resistance to an antitumor antibiotic: a DNA glycosylase triggers the base-excision repair system in yatakemycin biosynthesis. Angew Chem Int Ed Engl 51, 10532–10536.

Yan, Y., Liu, Q., Zang, X., Yuan, S., Bat-Erdene, U., Nguyen, C., Gan, J., Zhou, J., Jacobsen, S.E., and Tang, Y. (2018). Resistance-gene-directed discovery of a natural-product herbicide with a new mode of action. Nature 559, 415–418.

Zhang, M., Hou, X.F., Qi, L.H., Yin, Y., Li, Q., Pan, H.X., Chen, X.Y., and Tang, G.L. (2015). Biosynthesis of trioxacarcin revealing a different starter unit and complex tailoring steps for type II polyketide synthase. Chem Sci 6, 3440–3447.

Zhao, Q., He, Q., Ding, W., Tang, M., Kang, Q., Yu, Y., Deng, W., Zhang, Q., Fang, J., Tang, G., et al. (2008). Characterization of the azinomycin B biosynthetic gene cluster revealing a different iterative type I polyketide synthase for naphthoate biosynthesis. Chem Biol 15, 693–705.

Ziemert, N., Alanjary, M., and Weber, T. (2016). The evolution of genome mining in microbes -a review. Nat Prod Rep 33, 988–1005.

