## Supplemental Data for "Resistance-guided mining of bacterial genotoxins defines a family of DNA glycosylases"

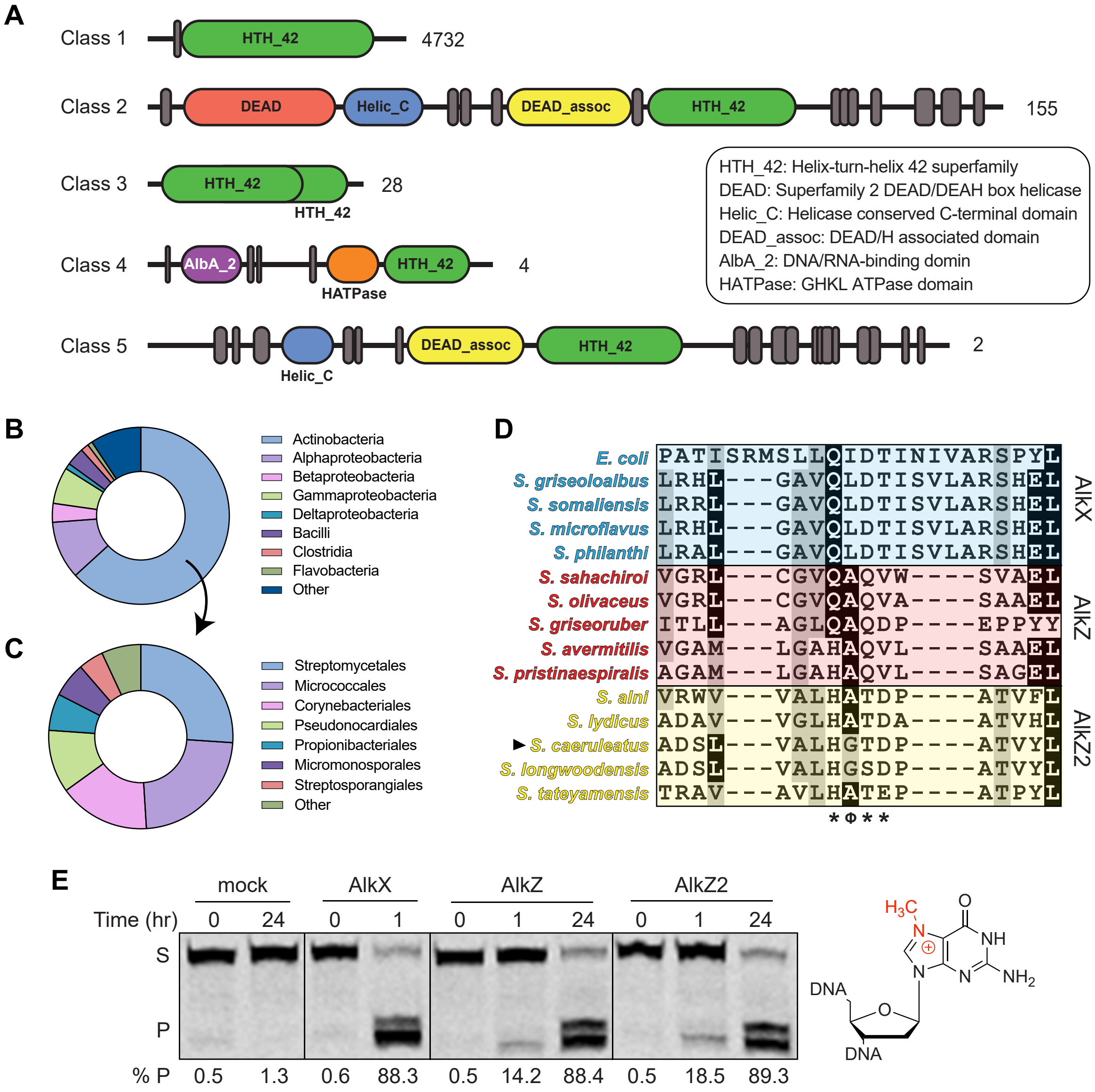


**Figure S1. HTH_42 superfamily taxonomy, phylogeny, and copy number analysis.** (**A**) Domain schematics for top 5 Pfam classes of HTH_42 superfamily proteins. Number of sequences for each organization is labeled to the right, along with the domain key. Dark vertical lines represent predicted unstructured regions. (**B**) Taxonomic distribution of HTH_42 proteins from prokaryotes (Class, 4,797 sequences). (**C**) Taxonomic distribution of HTH_42 proteins from Actinobacteria (Order, 3,033 sequences). (**D**) Sequence alignment of AlkX (blue) AlkZ (red), and AlkZ2 (yellow) proteins in the region surrounding the catalytic motif (asterisks). *Streptomyces caeruleatus* AlkZ2 (triangle) was used to test for glycosylase activity in panel E. (**E**) Denaturing PAGE of 5′-Cy5 labeled d7mG-DNA substrate (S) and nicked AP-DNA product (P) after treatment with enzyme or buffer (mock). AP-DNA resulting from glycosylase activity was nicked by treatment with 0.1 M NaOH to produce β,δ-elimination products, which are quantified below the gel.

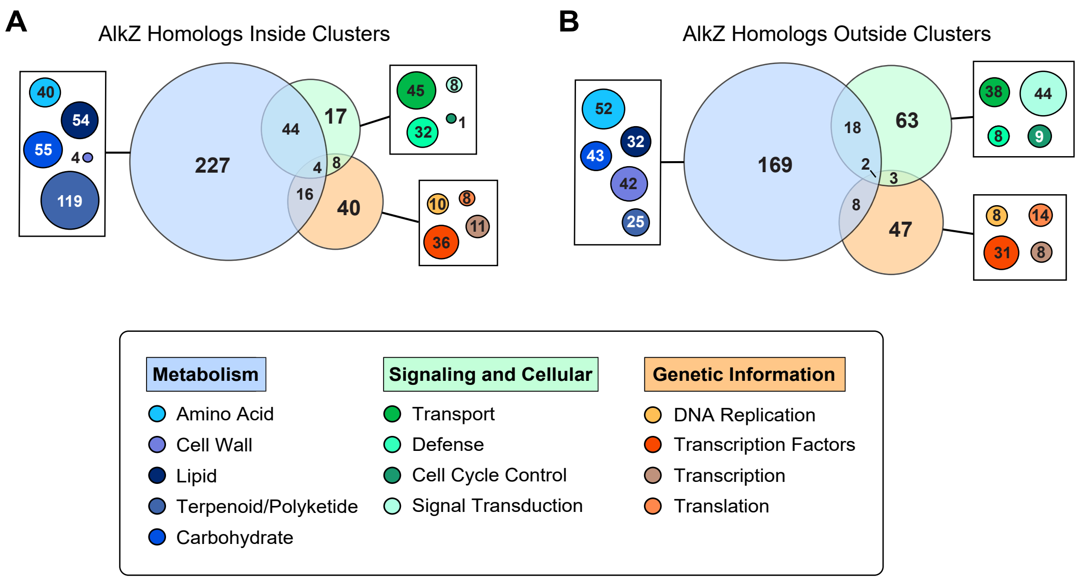


**Figure S2. Nearest neighbor analysis of AlkZ homologs.** Gene ontology (GO) analysis for nearest AlkZ neighbors (± 5 open reading frames) inside (**A**) and outside (**B**) BGCs. Venn diagrams depict the number of neighbors involved in metabolism (blue), signaling and cell function (green), and processing of genetic information (orange). The boxes represent subdivisions of each of the three functions, colored with respect to the key below. Uncharacterized/hypothetical proteins (40 inside, 90 outside) that could not be identified by homology are not included in these data. Full GO term analysis can be found in SI Tables S6-S11.

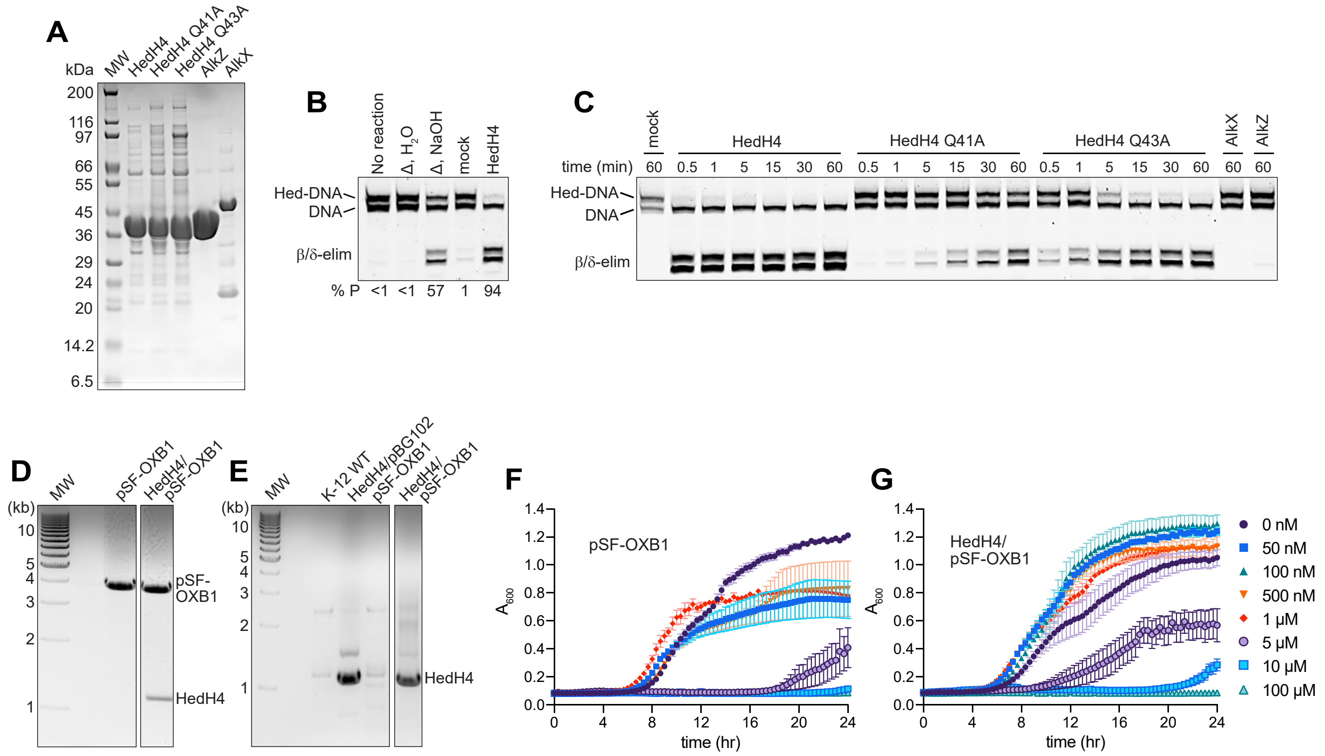


**Figure S3. HedH4 biochemistry and cellular resistance.** (**A**) Coomassie-stained SDS-PAGE of purified HedH4, *S. sahachiroi* AlkZ, and *E. coli* AlkX proteins. MW, molecular weight standards. Calculated protein molecular weights are 40.8 kDa (HedH4), 41.2 kDa (AlkZ), and 47.7 kDa (AlkX). (**B**) Thermal and enzyme-catalyzed depurination of Hed-DNA adducts. Denaturing PAGE of 5'-Cy5-labeled Hed-DNA olidodeoxynucleotide substrate and β- and δ-elimination products formed from hydroxide treatment of the abasic site generated from hydrolysis of the Hed-deoxyguanosine N-glycosidic bond. Formation of Hed-DNA goes to ~50% completion under our reaction conditions. Lane 1, Hed-DNA; lanes 2-3, Hed-DNA heated to 95°C for 5 min followed by treatment with either water or NaOH; lanes 4-5, Hed-DNA treated with either buffer (mock) or 1 μM HedH4 for 1 hour at 25°C, followed by NaOH. (**C**) Denaturing PAGE of hedamycin excision by HedH4 wild-type and catalytic mutants Q41A and Q43A, *E. coli* AlkX, and *S. sahachiroi* AlkZ. Mock, reaction with buffer alone. Quantification of this gel and the replicates are in Fig. 4D. (**D**,**E**) Verification of HedH4 cloning. (**D**) 1% agarose gel of analytical restriction digest of empty pSF-OXB1 and HedH4/pSF-OXB1 using NcoI-HF and XbaI restriction enzymes. Calculated molecular weights for pSF-OXB1 and HedH4 are 3.9 kb and 1.1 kb, respectively. (**E**) 1% agarose gel of colony PCR of HedH4 transformants in *E. coli* using HedH4 NcoI and XbaI primers (Table S1). Wild-type *E. coli* K-12 served as the negative control, while the protein expression vector HedH4/pBG102 served as a positive control. (**F**,**G**) Growth curves for *E. coli* K-12 containing either pSF-OXB1 (**F**) or HedH4/pSF-OXB1 (**G**) grown in LB/Kan media supplemented with increasing concentrations of hedamycin. Values are mean ± SD (n=3).

### Supplementary Tables

**Table S1. List of AlkX/AlkZ homologs by organism.** List of all *Streptomyces* AlkX, AlkZ, or AlkZ2 homologs in this study, along with the GenBank/RefSeq genome/assembly ID for each organism. Homologs are alphabetized by organism.

**Table S2. Proximity of AlkX/AlkZ homologs to biosynthetic gene clusters.** Results from AlkX/AlkZ BGC proximity analysis organized by their distances to the nearest antiSMASH-predicted cluster in the species' genome. Nearest cluster upstream (5'/(-)) and/or downstream (3'/(+)) is recorded with the cluster ID, and the most related cluster BLAST hit is denoted with the % gene similarity. *No BGC identified* denotes the cluster BLAST could not find a similar cluster which compares to the hit by homology.

**Table S3. AlkZ homologs found in uncharacterized biosynthetic gene clusters.** Information for AlkZ homologs in uncharacterized BGCs. *%I/S to AlkZ* (column C) is the percent identity or similarity to *S. sahachiroi* AlkZ. *Cluster BLAST* (column E) is the most similar BGC as determined by cluster BLAST analysis (*% similarity* is the percentage of genes in uncharacterized BGC that have homology to genes in the known similar BGC).

**Table S4. AlkZ homologs found in characterized biosynthetic gene clusters.** Information for AlkZ homologs in known BGCs. *%I/S to AlkZ* (column D) is the percent identity or similarity to *S. sahachiroi* AlkZ.

**Table S5. Oligodeoxynucleotides used in this study.** All oligos were dissolved in TE buffer (10 mM Tris•HCl pH 8.0, 1 mM EDTA pH 8.0) to 200 μM, and the DNA was stored at -20°C (stored in the dark for the Cy5-Hed oligo). The underlined nucleotide in the 7mG_Top and Hed_Top oligo is the site of the *N*7-alkylguanine lesion. PCR was performed with a primer concentration of 500 nM.

**Table S6. Nearest AlkZ neighbor GO term analysis for -5 ORFs within BGCs.** Nearest 5 open reading frames (ORFs) upstream (3' → 5') of AlkZ homologs predicted to be within BGCs. ORFs are listed with their GenBank/RefSeq ID and biological pathway and molecular GO terms as determined by NCBI, GhostKOALA, and eggNOG databases. Empty cells mean no GO terms could be assigned to these proteins through homology search.

**Table S7. Nearest AlkZ neighbor GO term analysis for +5 ORFs within BGCs.** Nearest 5 open reading frames (ORFs) downstream (5' → 3') of AlkZ homologs predicted to be within BGCs. ORFs are listed with their GenBank/RefSeq ID and biological pathway and molecular GO terms as determined by NCBI, GhostKOALA, and eggNOG databases. Empty cells mean no GO terms could be assigned to these proteins through homology search.

**Table S8. Nearest AlkZ neighbor GO term analysis for -5 ORFs outside BGCs.** Nearest 5 open reading frames (ORFs) upstream (3' → 5') of AlkZ homologs assigned to be outside BGCs. ORFs are listed with their GenBank/RefSeq ID and biological pathway and molecular GO terms as determined by NCBI, GhostKOALA, and eggNOG databases. Empty cells mean no GO terms could be assigned to these proteins through homology search.

**Table S9. Nearest AlkZ neighbor GO term analysis for +5 ORFs outside BGCs.** Nearest 5 open reading frames (ORFs) downstream (5' → 3') of AlkZ homologs assigned to be outside BGCs. ORFs are listed with their GenBank/RefSeq ID and biological pathway and molecular GO terms as determined by NCBI, GhostKOALA, and eggNOG databases. Empty cells mean no GO terms could be assigned to these proteins through homology search.

**Table S10. Nearest AlkX neighbor GO term analysis for -5 ORFs outside BGCs.** Nearest 5 open reading frames (ORFs) upstream (3' → 5') of AlkX homologs assigned to be outside BGCs. ORFs are listed with their GenBank/RefSeq ID and biological pathway and molecular GO terms as determined by NCBI, GhostKOALA, and eggNOG databases. Empty cells mean no GO terms could be assigned to these proteins through homology search.

**Table S11. Nearest AlkX neighbor GO term analysis for +5 ORFs outside BGCs.** Nearest 5 open reading frames (ORFs) downstream (5' → 3') of AlkX homologs assigned to be outside BGCs. ORFs are listed with their GenBank/RefSeq ID and biological pathway and molecular GO terms as determined by NCBI, GhostKOALA, and eggNOG databases. Empty cells mean no GO terms could be assigned to these proteins through homology search.
